# MicroRNA-27a is essential for bone remodeling by modulating p62-mediated osteoclast signaling

**DOI:** 10.1101/2022.06.13.495939

**Authors:** Shumin Wang, Eri O Maruyama, John Martinez, Justin Lopes, Trunee Hsu, Wencheng Wu, Wei Hsu, Takamitsu Maruyama

## Abstract

The ability to simultaneously modulate a set of genes for lineage-specific development has made microRNA an ideal master regulator for organogenesis. However, most microRNA deletions do not exhibit obvious phenotypic defects possibly due to functional redundancy. MicroRNAs are known to regulate skeletal lineages as the loss of their maturation enzyme Dicer impairs bone remodeling processes. Therefore, it is important to identify specific microRNA essential for bone homeostasis. We report the loss of miR-27a causing severe osteoporosis in mice. MiR-27a affects osteoclast-mediated bone resorption but not osteoblast-mediated bone formation during skeletal remodeling. Gene profiling and bioinformatics further identify the specific targets of miR-27a in osteoclast cells. MiR-27a exerts its effects on osteoclast differentiation through modulation of Squstm1/p62 whose mutations have been linked to Paget’s disease of bone. Our findings reveal a new miR-27a-p62 axis necessary and sufficient to mediate osteoclast differentiation and highlight a therapeutic implication for osteoporosis.

## Introduction

MicroRNA (miRNA) is a small non-coding RNA, base-pairing with complementary sequences of messenger (mRNA) to control gene expression at post-transcriptional and translational levels ^1, 2^. In animals, miRNA recognizes the 3’ untranslated region of their targets via a small stretch of seed sequences. A single miRNA may simultaneously affect the expression of hundreds of genes ^3, 4^. Because of its potential to modulate the same biological process at various steps, miRNA has been postulated to function as a master regulator for organogenesis ^5, 6^, similar to the transcription factor capable of turning on a set of genes for lineage-specific development. However, possibly due to functional redundancy, most of the miRNA deletions do not cause phenotypic alterations ^7, 8^. With only limited evidence ^9, 10^, it has been difficult to prove this concept.

The skeleton is constantly remodeled to maintain a healthy structure after its formation ^11^. This lifelong process is called bone remodeling in which old bone is removed from the skeleton, followed by replaced with new bone. Therefore, a balance of osteoclast-mediated bone resorption and osteoblast-mediated bone formation is essential for bone metabolism ^11^. Dysregulation of bone remodeling causes metabolic disorders, e.g. osteopetrosis, osteoporosis, and Paget’s disease ^12^. The current treatments have major limitations leading to the exploration of new therapeutic strategies. Studies of Dicer, an RNase III endonuclease involved in the maturation of miRNAs, suggest their importance in the development of the skeletal lineages during bone remodeling ^13, 14^. However, the specific miRNA(s) required for differentiation of osteoclasts and osteoblasts remains largely unclear. The miR-23a cluster consists of miR-23a, miR-27a, and miR-24-2. Aberrant regulation of miR-23a and miR-27a has been associated with osteoporotic patients and increased bone fracture risks ^15–17^. The effect of miR-23a and miR-27a on the differentiation of osteoblast or osteoclast cells has been shown by in vitro overexpression studies ^16, 18, 19^. Interference of miR-23a or miR-27a by the use of inhibitor/sponge has implied their role in osteoblast and osteoclast cells ^18, 20^. However, due to cross-reactivity of the RNA inhibitor among family members that share common seed motif ^21, 22^, the inhibitor assay may not truthfully reflect the function of the target miRNA. Therefore, the genetic loss-of-function study remains the most rigorous method for determining the endogenous function of miR-23a and miR-27a as well as testing the removal of miR-23a∼27a sufficient to cause bone loss.

We have performed mouse genetic analyses to definitively assess the requirement of miR-23a and miR-27a for skeletogenesis and homeostasis. Surprisingly, the skeletal phenotypes developed in newly established loss-of-function mouse models reveal findings different from previous reports based on the gain-of-function analyses. Severe loss of bone mass developed in mice with the deletion of miR-23a∼27a or miR-27a. MiR-23a∼27a is dispensable for osteoblast-mediated bone formation. However, compelling evidence support that miR-27a is essential for osteoclastogenesis and osteoclast-mediated bone resorption during bone remodeling. Gene expression profiling and bioinformatics analyses further identify osteoclast-specific targets of miR-27a. We demonstrated that miR-27a exerts its effects on osteoclast differentiation through modulation of a new and essential target Squstm1/p62. MiR-27a is necessary and sufficient to mediate osteoclast differentiation, and as a biomarker and therapeutic target for osteoporosis.

## Results

### The loss of miR23a∼27a in mice causes low bone mass phenotypes

To determine the requirement of miR-23a∼27a for skeletal development and maintenance, we created a new mouse model with the deletion of miR-27a and miR-23a. The CRISPR/Cas9 gene edition method was used to establish the mouse strain carrying the ΔmiR-23a∼27a allele (Fig. 1A). PCR analysis of the miR-23a cluster demonstrated the sgRNA-mediated deletion and the reduction of 500 bp in the wild-type to 303 bp in the mutants (Fig. 1B). Sequencing analysis also confirmed the expected genome editing (data not shown). Next, we tested if the deletions affect the expression of other microRNA molecules, generated from the same RNA precursor, within the same cluster. Semi-quantitative RT-PCR analysis revealed that only miR-23a∼27a are disrupted in the ΔmiR-23a∼27a mutants (Fig. 1C). These results indicated our success in establishing mouse models deficient for miR-23a∼27a.

**Fig. 1.**
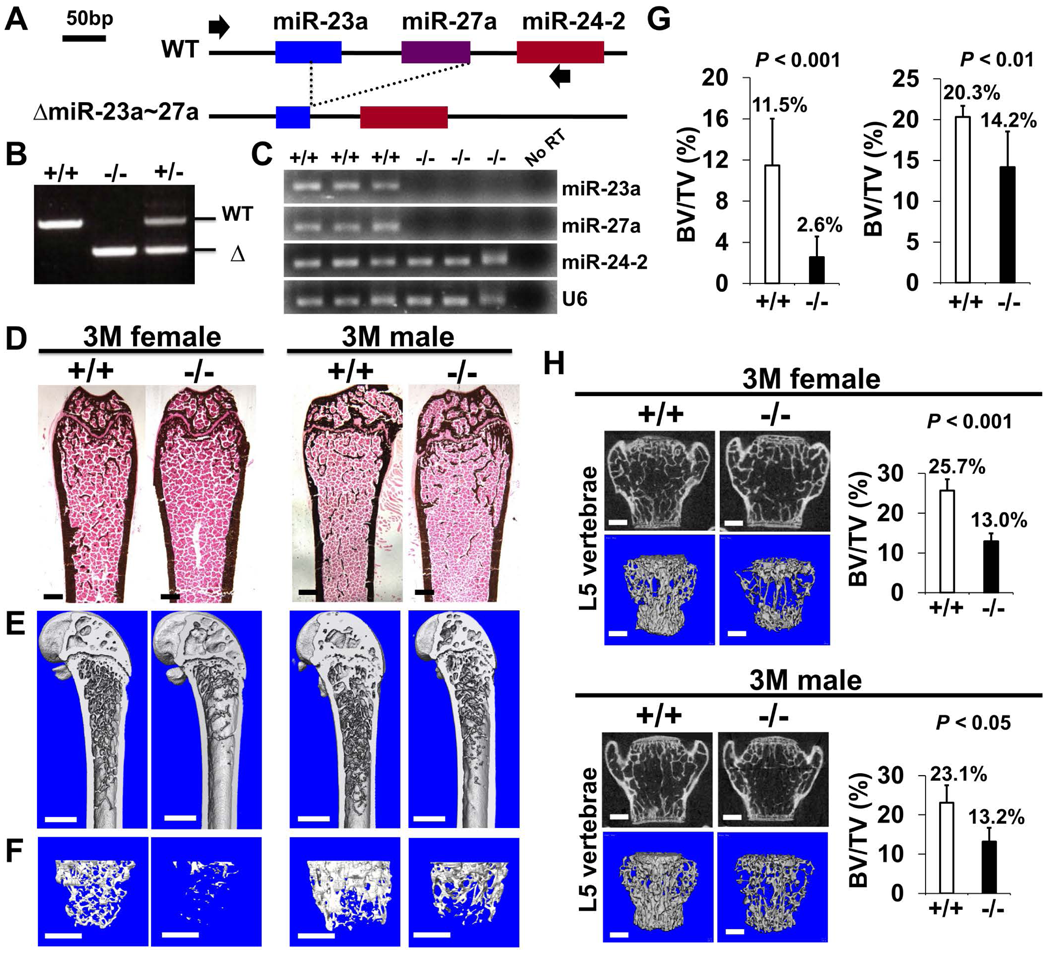
Low bone mass phenotypes in mice deficient for miR-23a∼27a. (A) Diagrams illustrate the miR-23a cluster, consisting of miR-23a, miR-27a, and miR-24-2 (WT), and the creation of mouse strains deficient for miR-23a∼27a (ΔmiR-23a∼27a) by CRISPR/Cas9 genome editing. Broken lines and arrows indicate the deleted genomic regions and primers used for PCR genotyping analysis, respectively. (B) PCR analysis examines the miR-23a cluster for genotyping the wild-type (+/+), heterozygous (+/-) and homozygous (-/-) for miR-23a∼27a mice. The mutant (Δ) alleles with deletion of miR-23a∼27a result in the generation of shorter PCR products. (C) RT-PCR analysis of the miR-23a, miR-27a, and miR-24-2 RNAs reveals the disruption of specific microRNA(s) in the mutants. The analysis of small noncoding RNA U6 is used as an internal control. Femurs of the 3-month-old (3M) wild-type (+/+) and mutant (-/-) males and females were analyzed by μCT scanning (E-F), followed by sectioning and von Kossa staining (D). Reconstructed μCT images of the distal femur (E) and femoral metaphysis (F) were subject to quantitative analysis for trabecular bone volume (G). (H) Spines of the 3-month-old (3M) ΔmiR-23a∼27a males and females were analyzed by μCT scanning. Images show the μCT scanned wild-type (+/+) and mutant (-/-) L5 vertebrae (top) and 3D rendered trabecular bone (bottom). Quantitative analyses of trabecular bone volume per total volume in the femurs and vertebrates are shown in graphs (BV/TV, n=3, mean ± SD; student t-test). Images (D-F, H) are representatives of three independent experiments. Scale bars, 500 µm (D-F, H).

Mice heterozygous for ΔmiR-23a∼27a are viable and fertile. Intercross between the heterozygotes successfully obtained the homozygous mutants without any noticeable skeletal deformity, suggesting that miR-23a∼27a are not required for the developmental processes (Fig. S1). Next, we examined if their deletions affect the homeostatic maintenance of the bone in adults. At 3 months, von Kossa staining and three-dimensional (3D) micro-computed tomography (µCT) analyses of the ΔmiR-23a∼27a femurs revealed significant loss of the trabecular bone volume in both sexes (Fig. 1D-G; BV/TV, n=3, mean ± SD; student t-test). Much more severe osteoporotic defects were detected in the 7-month-old mutant females of ΔmiR-23a∼27a (Fig. S2; BV/TV, n=3, mean + SD; student t-test). However, cortical bone thickness does not seem to be affected by the deletion of miR-23a∼27a (Fig. S3). Next, we examined if similar bone loss phenotypes can be detected in the vertebrae where age-related changes in the trabecular architecture are minimal. Therefore, we examined the vertebrae of ΔmiR-23a∼27a and identified drastic reductions in vertebral bone mass associated with the mutations (Fig. 1H). These data demonstrated that miR-23a∼27a is required for homeostatic maintenance of the bone.

### Osteoblast-mediated bone formation is not affected by the loss of miR-23a∼27a

Proper maintenance of the skeleton requires balanced bone formation and resorption during bone remodeling. The bone loss phenotypes caused by the deletion of miR-23a∼27a are likely to be associated with an imbalanced bone formation and resorption mediated by osteoblasts and osteoclasts, respectively. Therefore, we examined if the bone formation and osteoblast activities are affected by the loss of miR-23a∼27a.

New bone formation was analyzed by double labeling with alizarin red and calcein at 3 months. Quantitative analyses did not reveal a significant difference in bone formation rate per unit of bone surface (BFR/BS) caused by the mutation (Fig. S4A). In addition, the numbers of osteoblast cells positive for type 1 collagen (Col1) and Osteopontin (OPN) lining the trabecular bone surface remain comparable between the wild-type and homozygous littermates (Fig. S4B), indicating that osteoblast-mediated bone formation is not affected by the miR-23a∼27a deletion. The results suggested that miR-27a and miR-23a are not required for osteoblastogenesis and osteoblasts-mediated bone formation.

### MiR-23a∼27a regulates osteoclast differentiation

To determine if the loss of miR-27a affects bone resorption, we first examined osteoclast number by tartrate-resistant acid phosphatase (TRAP) staining. An increase of TRAP+ osteoclasts was detected in the 3-month-old ΔmiR-23a∼27a males and females (Fig. 2). When the number of TARP+ osteoclast cells in the total bone area (N.Oc/T.Ar), the ratio of TRAP+ bone surface (Oc.S/BS), and the number of TRAP+ osteoclast cells lining the bone surface (N.Oc/BS) were measured, we found that these parameters associated with bone resorption are significantly elevated in the mutants (Fig. 2; n=5, mean ± SD; student t-test). In addition, there is a ∼3-fold increase in the number of Cathepsin K-expressing osteoclast cells lining the trabecular bone surface (Fig. 2; n=5, mean ± SD; student t-test). These results support that the loss of miR-23a∼27a stimulates osteoclastogenesis, leading to an elevation of bone resorption.

**Fig. 2.**
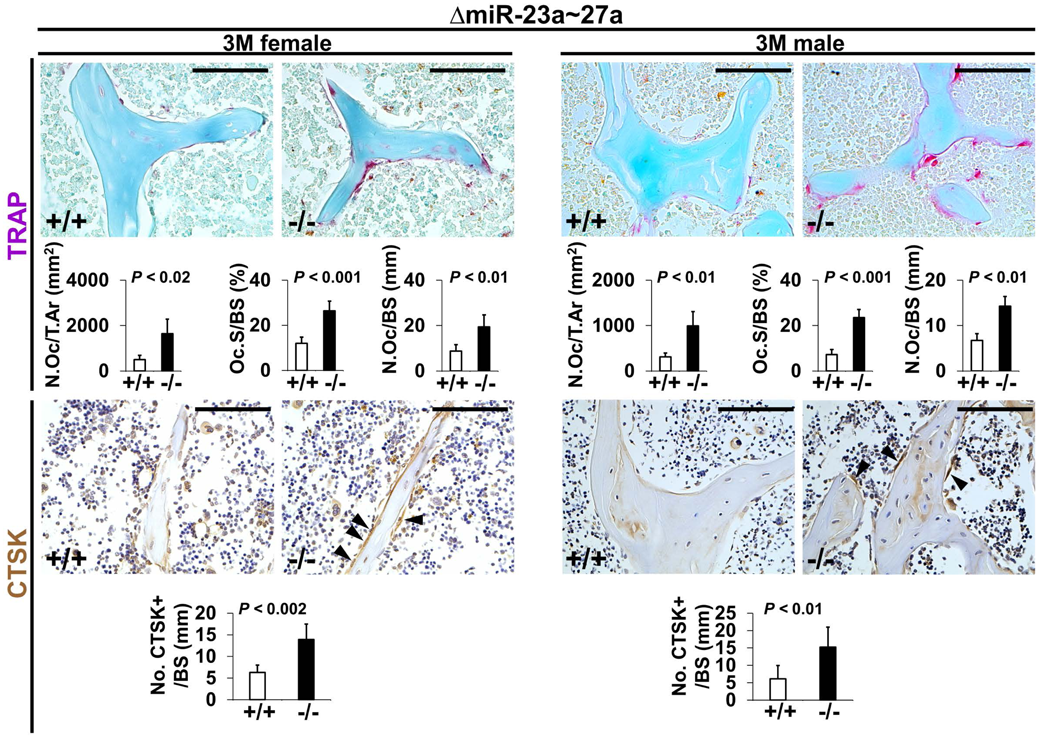
Increased number of osteoclast cells in the ΔmiR-23a∼27a mice. Sections of the 3-month-old (3M) ΔmiR-23a∼27a males and females were analyzed by tartrate-resistant acid phosphatase (TRAP) staining and immunostaining of Cathepsin K (CTSK). Graphs show quantitative analyses of positively stained cells in the wild-type (+/+) and mutant (-/-) distal femurs (No. of cell+/BS, n=5, mean ± SD; student t-test). Histomorphometric parameters of bone resorption are evaluated by number of osteoclast/bone area (N.OC/T.Ar), osteoclast surface/bone surface (OC.S/BS), osteoclast number/bone surface (N.OC/BS, n=5, mean ± SD; student t-test). Images are representatives of five independent experiments. Scale bars, 100 µm.

Next, to determine the role of miR-23a∼27a in osteoclastogenesis, we analyzed cell populations associated with the differentiation of the osteoclast cells. During hematopoiesis, a common myeloid progenitor gives rise to monocytes that are precursors of several cell types, including dendritic cells, macrophages, and osteoclasts ^23^. Osteoclast precursors are known to derive from a monocyte population positive for CD11b and negative for Gr-1 ^24^. Therefore, we examined CD11b+ and Gr-1– monocytes to see if the miR-23a∼27a deletion affects the osteoclast precursor population. FACS analysis revealed that the CD11b+ and Gr-1– population was not affected in the ΔmiR-23a∼27a bone marrow of both male and female mice (Fig. 3A, B). The CD11b+/CD11c+ dendritic cell population, derived from the monocytes, was also unaffected by the miR-23a∼27a deletion (Fig. 3A, B). Although the precursors are not affected, miR-23a∼27a may play a role in osteoclast differentiation.

**Fig. 3.**
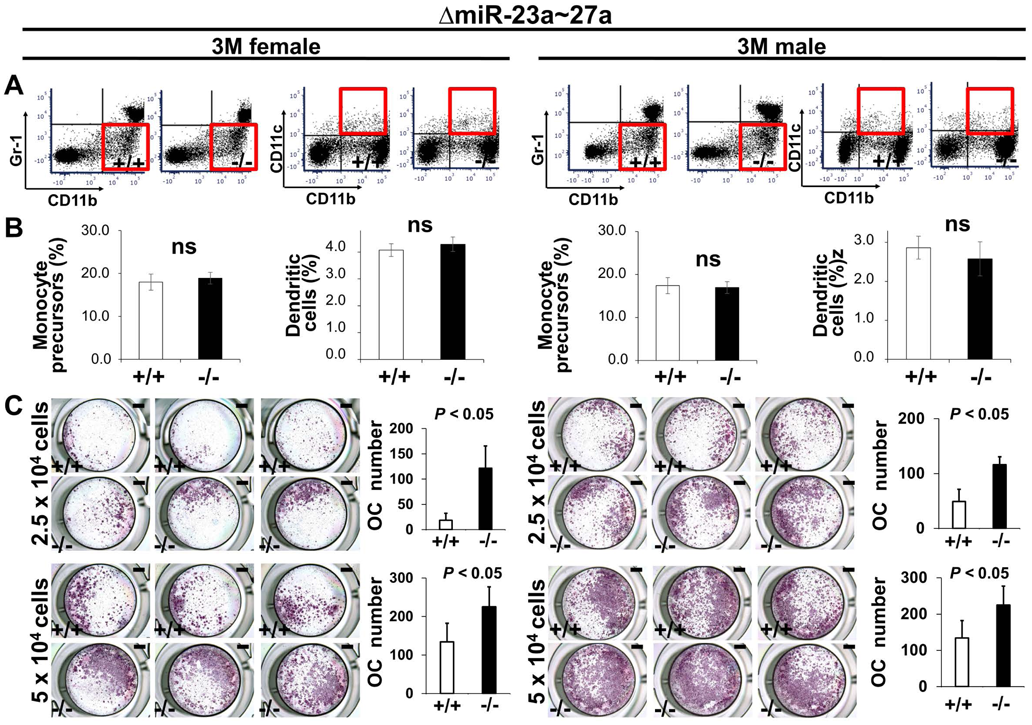
MiR-23a∼27a regulates osteoclast differentiation. (A) FACS analysis examines the CD11b+/Gr-1- and CD11b+/CD11c+ populations for monocyte precursors and dendritic cells, respectively. Images are representatives of three independent experiments. (B) The bone marrow of 3-month-old (3M) wild-type (+/+) and ΔmiR-23a∼27a (-/-) males and females show no significant difference (ns; n = 3, mean ± SD; student t-test). (C) Cells isolated from the bone marrow were induced for osteoclast (OC) differentiation with RANLK and MCSF for 5 days (top, 2.5×10^4^ cells/well and bottom, 5×10^4^ cells/well). The number of OC cells positive for TRAP staining is significantly enhanced in the mutant cultures (n=5, mean ± SD; student t-test). Scale bars, 1 mm (C).

To test osteoclast differentiation affected by the loss of miR-23a∼27a, an ex vivo analysis was performed with cultures of cells seeding at two different densities. Cells isolated from the bone marrow were cultured in the presence of M-CSF to obtain bone marrow-derived macrophages (BMMs), followed by differentiation into osteoclasts with the treatment of RANKL. TRAP staining was then used to assess the extent of osteoclast differentiation. The number of TRAP+ cells was significantly increased by the loss of miR-23a∼27a (Fig. 3C; n=5, mean ± SD; student t-test).

### MiR-27a is an essential regulator for osteoclast-mediated skeletal remodeling

Using the miRPath Reverse-Search module, we searched and ranked miRNAs whose targets are accumulated in osteoclast differentiation-related genes based on the enrichment of the targets in the Kyoto Encyclopedia of Genes ad Genomics (KEGG: mmu04380) ^25, 26^. Among them, miR-27a was predicted as the top 3 candidate to regulate osteoclast differentiation (Fig. S5, p < 2.0 x 10^-71^). Its sister gene miR-27b contains the same seed sequences also ranked in the top 3. However, miR-23a had a lower estimated rank suggesting that miR-27a alone may be sufficient to exert osteoclast regulation (Fig. S5, p < 4.9 x 10^-29^). To test this hypothesis, we created another mouse strain with the deletion of only miR-27a using the CRISPR/Cas9 genome editing (Fig. 4A). PCR analysis revealed the sgRNA-mediated deletion causes the reduction of 500 bp in the wild-type to 474 bp in the mutants (Fig. 4B). Sequencing of the PCR products confirmed the genomic deletion (data not shown). Next, semi-quantitative RT-PCR analysis indicated that only miR-27a are disrupted in the ΔmiR-27a mutants, suggesting the deletions does not affect the expression of other microRNA generated from the same RNA precursor (Fig. 4C). These results demonstrated our success in establishing mouse models deficient for miR-27a.

**Fig. 4.**
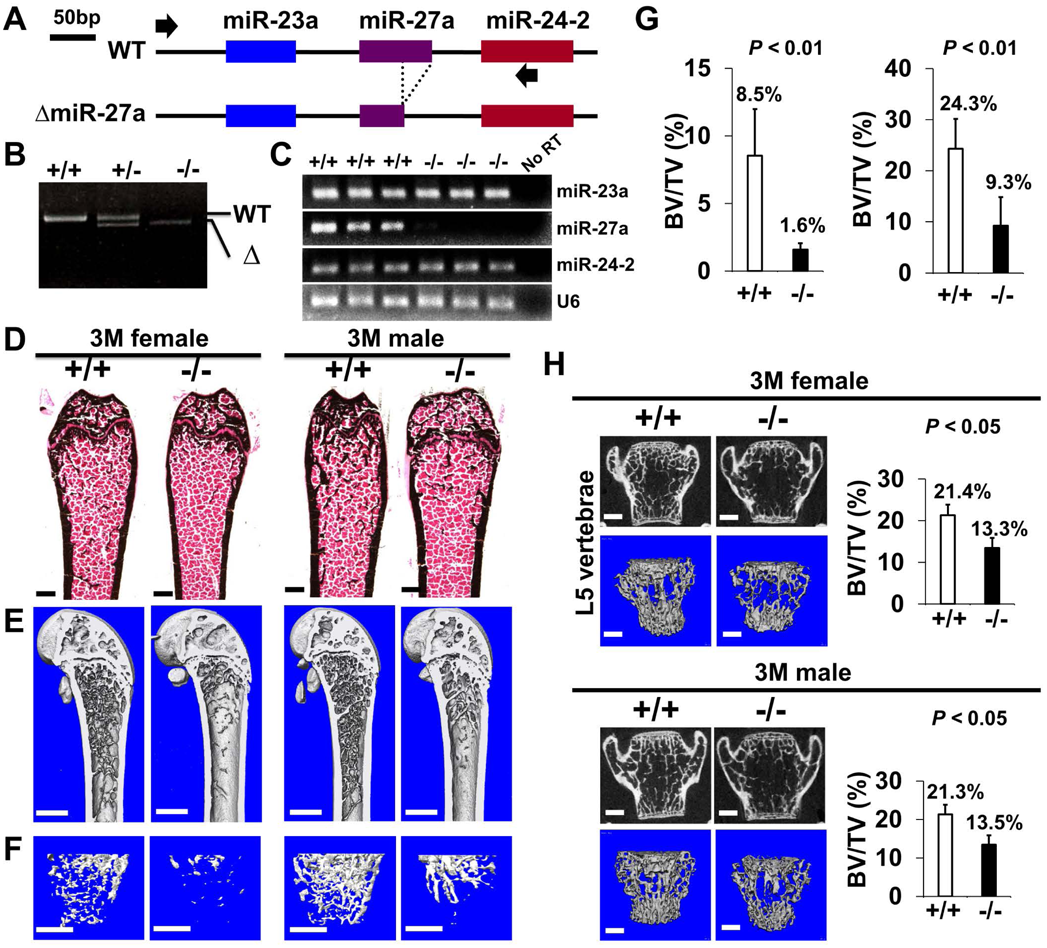
The loss of miR-27a alone causes osteoporotic defects. (A) Diagrams illustrate the miR-23a cluster (WT), and the creation of mouse strains deficient for miR-27a (ΔmiR-27a) by CRISPR/Cas9 genome editing. Broken lines and arrows indicate the deleted genomic regions and primers used for PCR genotyping analysis, respectively. (B) PCR-based genotyping identifies the wild-type (+/+), heterozygous (+/-) and homozygous (-/-) for miR-27a mice showing the mutant (Δ) alleles with deletion of miR-27a result in the generation of shorter PCR products. (C) RT-PCR analysis reveals the disruption of miR-27a but not miR-23a and miR-24-2 RNAs in ΔmiR-27a mutants. The analysis of small noncoding RNA U6 is used as an internal control. Femurs of the 3-month-old (3M) wild-type (+/+) and mutant (-/-) males and females were analyzed by μCT scanning (E-F), followed by sectioning and von Kossa staining (D). The 3D rendered μCT images of the distal femur (E) and femoral metaphysis (F) were subject to quantitative analysis for trabecular bone volume (G). (H) Spines of the 3-month-old (3M) ΔmiR-27a males and females were analyzed by μCT scanning. Images show the μCT scanned wild-type (+/+) and mutant (-/-) L5 vertebrae (top) and 3D rendered trabecular bone (bottom). Quantitative analyses of trabecular bone volume per total volume in the femurs and vertebrates are shown in graphs (BV/TV, n=3, mean ± SD; student t-test). Images (D-F, H) are representatives of three independent experiments. Scale bars, 500 µm (D-F, H).

Mice heterozygous and homozygous for ΔmiR-27a are viable and fertile similar to the miR-23a∼27a deletion. As anticipated, there was no noticeable skeletal deformity associated with the loss of miR-27a suggesting its dispensable role in the developmental processes. However, von Kossa staining and 3D micro-computed tomography (µCT) analyses of the 3-month-old male and female femurs of ΔmiR-27a revealed significant loss of the trabecular bone volume (Fig. 4D-G; BV/TV, n=3, mean ± SD; student t-test) while cortical bone thickness was not affected (Fig. S6). Drastic bone loss phenotypes can also be detected in the ΔmiR-27a vertebrae where age-related changes in the trabecular architecture are minimal (Fig. 4H). These data demonstrated the essential role of miR-27a in skeletal remodeling.

Using double labeling with alizarin red and calcein for quantitative analyses, we did not reveal a significant difference in bone formation rate per unit of bone surface (BFR/BS) (Fig. S7A), and Col1+ and OPN+ osteoblast cells lining the trabecular bone surface (Fig. S7B) at 3 months, suggesting that osteoblast-mediated bone formation is not affected by the miR-27a deletion. However, we detected a significant increase of TRAP+ and Cathepsin K-expressing osteoclast cells lining the trabecular bone surface in the 3-month-old ΔmiR-27a males and females (Fig. 5A and Fig. S8; n=5, mean ± SD; student t-test), supporting that the loss of miR-27a stimulates osteoclastogenesis, leading to elevated bone resorption. While the CD11b+/Gr-1– and CD11b+/CD11c+ osteoclast precursor populations were not affected by the miR-27a deletion (Fig. S9A, B), osteoclast differentiation was significantly increased by the loss of miR-27a (Fig. 5B; n=5, mean ± SD; student t-test). Furthermore, the loss of a single miR-27a recapitulates the osteoporotic phenotypes caused by the double deletion of miR-23a and miR-27a, suggesting that miR-27a is responsible for skeletal maintenance through the modulation of bone remodeling processes. The results suggested that miR-27a functions as a negative regulator in osteoclast differentiation. To test this possibility, we overexpressed miR-27a in cells undergoing osteoclast differentiation. High levels of miR-27a significantly reduce the number of differentiated osteoclast cells (Fig. 5C). Our findings demonstrated that miR-27a is necessary and sufficient to modulate osteoclast differentiation. Osteoclastogenesis mediated by miR-27a is essential for bone remodeling and homeostasis.

**Fig. 5.**
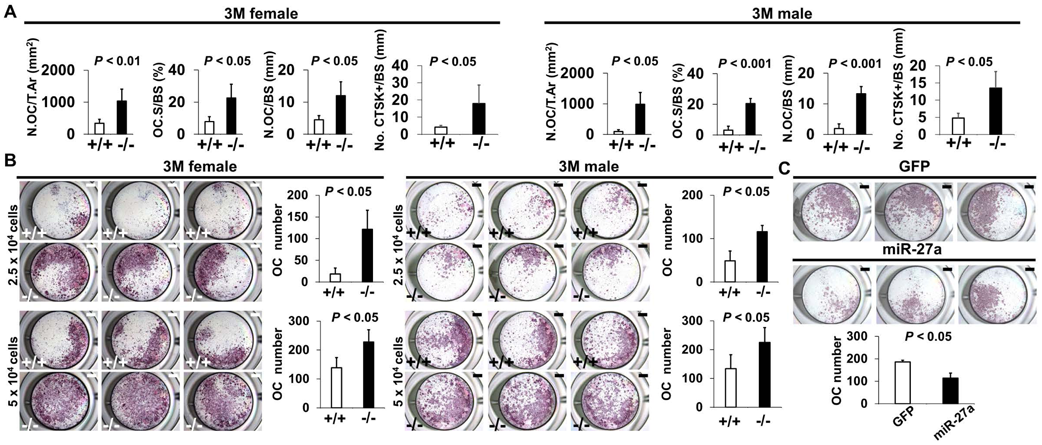
MiR-27a is necessary and sufficient to repress osteoclast differentiation. (A) Graphs show quantitative analyses of TRAP+ and CTST+ cells in the wild-type (+/+) and mutant (-/-) distal femurs detected in the stained section of the 3-month-old (3M) ΔmiR-27a males and females (n=5, mean ± SD; student t-test). Histomorphometric parameters of bone resorption are evaluated by number of osteoclast/bone area (N.OC/T.Ar), osteoclast surface/bone surface (OC.S/BS), osteoclast number/bone surface (N.OC/BS), CTSK+ cells/bone surface (No. CTSK+/BS). (B) Cells isolated from the wild-type (+/+) and mutant (-/-) bone marrows were induced for osteoclast (OC) differentiation with RANLK and MCSF for 5 days (top, 2.5×10^4^ cells/well and bottom, 5×10^4^ cells/well). Graphs indicate the number of TRAP+ OC cells (n=5, mean ± SD; student t-test). (C) Cells isolated from the bone marrow were seeded (5×10^4^ cells/well) and induced for OC differentiation using RANLK and MCSF for 5 days with the lentivirus-mediated expression of GFP (control) or miR-27a. The TRAP+ OC cell number is significantly decreased in the miR-27 overexpression cultures (n=3, p < 0.05, mean ± SD; student t-test). Scale bars, 1 mm (B, C).

### MiR-27a regulates osteoclast differentiation through the modulation of p62

To elucidate the mechanism underlying OC differentiation regulated by miR-27a, we first used a bioinformatics approach to identify its potential targets (Fig. 6A). The TarBase computationally predicted 2312 target genes for miR-27a ^27^. Furthermore, there were 154 genes associated with osteoclast differentiation based on the Kyoto Encyclopedia of Genes ad Genomics (KEGG: mmu04380) ^25, 26^. The miRPath software further identified 26 targets overlapping with the osteoclast-related genes ^28^. Next, we examined the transcript level of these 26 targets in wild-type and ΔmiR-27a osteoclast cells. Quantitative RT-PCR analyses revealed that 5 of these targets, Snx10, Map2k7, Ctsk, Tgfbr1, and Sqstm1, are significantly up-regulated by the loss of miR-27a (Fig. 6B, p < 0.05, n = 3; two-sided student t-test). To test if these genes were the direct targets of miR-27a, we performed 3’UTR-reporter assays. The expression of miR-27a significantly downregulated the luciferase activity associated with the 3’UTR of Sqstm1, Tgfbr1, Snx10, Map2k7, but not Ctsk (Fig. 6C, *; p < 0.01, n=3, mean ± SD; two-sided student t-test). As Cathepsin K is a protease expressed in the mature osteoclast cells, its alteration at the transcript level is likely ascribed to indirect effects of increased osteoclastogenesis in ΔmiR-27a. The data indicated Sqstm1, Tgfbr1, Snx10, and Map2k7 as direct targets of miR-27a.

**Fig. 6.**
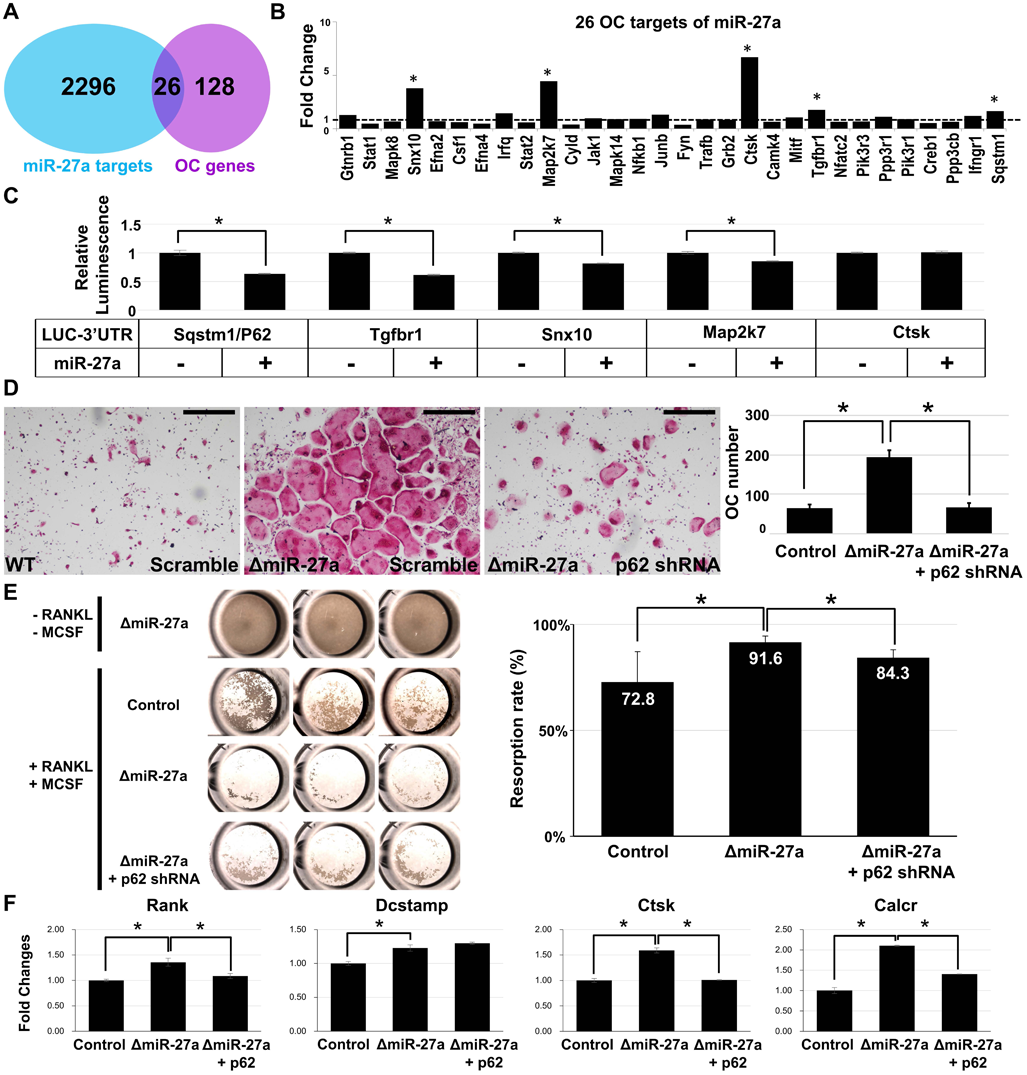
MiR-27a-dependent regulation of osteoclast differentiation is mediated through p62 modulation. (A) Venn diagrams illustrate our strategy to identify the OC differentiation-associated genes (KEGG: mmu04380) that are directly regulated by miR-27a. (B) The twenty-six potential targets are examined by quantitative RT-PCR analysis to detect the change of transcript levels in wild-type and ΔmiR-27a OC cells (n=3; *, *p*-value < 0.05, two-sided student’s t-test). (C) The 3’UTR-reporter assay examines five potential genes that are direct targets of miR-27a (n=3; *, p-value < 0.01, means ± SD, two-sided student’s t-test). (D) A functional study of Sqstm1 also known as p62 reveals the enhancement of OC differentiation caused by the loss of miR-27a is alleviated by lentiviral-shRNA-mediated knockdown. TRAP staining examines the number of mature OC cells in the control, ΔmiR-27a and ΔmiR-27a plus shRNA-mediated knockdown of p62 (ΔmiR-27a + p62 shRNA) cultures (n=3; *, p-value < 0.05, means ± SD, two-sided student’s t-test). (E) Pit assay examined OC cells-mediated bone resorption rate of control, ΔmiR-27a, ΔmiR-27a plus shRNA-mediated p62 knockdown 5 days after differentiation (n=3; *, p-value < 0.05, mean ± SD, two-sided student t-test). Osteoclast progenitors isolated from ΔmiR-27a mice were cultured without RANKL and MCSF as negative control (Top raw). (F) qRT-PCR examined the expression of osteoclast markers after 3 days (Rank and Dcstamp) or 7 days (Ctsk and Calcr) culture of the control, ΔmiR-27a, and ΔmiR-27a plus shRNA-mediated knockdown of p62 OC cells (n=3; *, p-value < 0.05, mean ± SD, student t-test). Scale bars, 500 µm.

Sqstm1 also known as p62 whose gain of function mutations were linked to Paget’s disease of bone with disruption of bone renewal cycle causing weakening and deformity ^29^. The deletion of p62 in mice also impaired osteoclast differentiation ^30^. Therefore, we performed a functional study to test the importance of the miR-27a-p62 regulatory axis during osteoblastogenesis. Cells isolated from the bone marrow were induced for osteoclast differentiation and the number of osteoclast cells positive for TRAP staining was counted to determine the outcome of the differentiation. The enhanced osteoclast differentiation in the ΔmiR-27a culture was significantly alleviated by the shRNA-mediated knockdown of p62 (Fig. 6D, p < 0.05, n=3, mean ± SD; two-sided student t-test). To assess osteoclast function, we first performed phalloidin staining to ensure actin ring formation was not affected by miR-27a deletion and p62 knockdown in OC cells (Fig. S10). Bone resorption pit assay then revealed enhanced bone resorption activity of ΔmiR-27a osteoclasts that can be alleviated by p62 knockdown (Fig. 6E, p < 0.05, n=3, mean ± SD; two-sided student t-test). Next, qRT-PCR of osteoclast markers was performed to decipher the regulatory process of miR-27a-mediated osteoclast differentiation. The deletion of miR-27a elevated the expression of Rank but this elevation could be alleviated by p62 knockdown (Fig. 6F, p < 0.05, n=3, mean ± SD; two-sided student t-test), indicating the importance of the miR-27a-p62 axis for RANKL signaling. The expression of Dcstamp, essential for cell-cell fusion during osteoclastogenesis, was increased in ΔmiR-27a osteoclasts (Fig. 6F, p < 0.05, n=3, mean ± SD; two-sided student t-test). However, the increased level of Dcstamp in ΔmiR-27a osteoclasts could not be affected by p62 knockdown, implying that miR-27a mediated regulation of osteoclast fusion is independent of p62. Mature osteoclast markers, e.g. Ctsk and Calcr, were significantly upregulated in ΔmiR-27a osteoclasts but reduced by p62 knockdown (Fig. 6F, p < 0.05, n=3, mean±SD; two-sided student t-test). The results demonstrated that miR-27a-dependent osteoclast differentiation is mediated through the regulation of p62. The miR-27a-p62 regulatory axis plays an important role in osteoclastogenesis during bone remodeling.

## Discussion

The dysregulation of miRNA has been implicated in osteoporosis in menopausal women. Among 851 miRNAs tested miR-27a is one of the most significant genes downregulated in postmenopausal osteoporosis patients ^17^. However, it’s not clear whether the alteration of miRNAs is the cause or consequence of the disease. Our genetic study presented here has demonstrated the loss of miR23a∼27a or miR27a results in significant bone loss. The findings suggest a single miRNA deficiency can lead to severe osteoporotic defects, indicating an essential role of miR-27a in bone remodeling. Because osteoporosis is caused by an imbalance of osteoblast-mediated bone formation and osteoclast-mediated bone resorption, the conventional knockout model is ideal to decipher the regulatory processes underlying miR-27a-dependent pathogenesis. Our data also suggest that miR-27a is dispensable for osteoblast differentiation and bone formation as its deletion does not affect the number of osteoblast cells and bone formation rates. The results do not agree with the previous gain-of-function study indicating an inhibitory role of miR-27a in osteoblastogenesis ^18^. Therefore, the association of osteoporosis with both upregulation and downregulation is possibly mediated through distinct mechanisms underlying the regulatory process of the miR-23a cluster.

This study provides compelling evidence to first demonstrate that miR-27a is essential for regulating the bone resorption process through modulation of osteoclast differentiation. The loss of miR-27a in mice leads to elevated numbers of osteoclast cells as well as increases in bone resorption activity. The inhibitory function of miR-27a on osteoclastogenesis is also in agreement with previous in vitro culture data showing its crucial downregulation among miRNAs associated with osteoclast differentiation ^31^. Furthermore, miR-23a upregulation has been detected in osteopetrosis patients ^32^. Although the number of osteoclast progenitors is comparable between the control and mutant, the deletion of miR-27a strongly accelerates the process of osteoclastogenesis. The isolated ΔmiR-27a exhibits a highly potent ability in differentiation and maturation, indicating cell-autonomous regulations of miR-27a in the osteoclast cell.

The gain-of-function mutations have linked p62 to the cause of Paget’s disease of bone – a genetic disorder characterized by aberrant osteoclastic activity ^29, 33^. The knockout of p62 in mice further supports its critical role in osteoclastogenesis ^29^. Our identification and characterization of p62 as a direct downstream regulator of miR-27a established a new osteoclast signaling axis. Not only osteoclast differentiation and maturation are suppressed by high levels of p62 but also its reduction can alleviate excessive osteoclastogenesis caused by the loss of miR-27a. Our findings suggest the miR-27a-p62 regulatory axis is necessary and sufficient to regulate bone remodeling through modulation of osteoclastogenesis. In addition, circulatory miRNAs were released in the cell-free form either bound with protein components or encapsulated with microvesicles. They are quite stable and found with variations in miRNA signature as biomarkers ^34^. Therefore, the identification of miR-27a as essential for bone remodeling promises its use as a biomarker for early detection of bone destruction-associated diseases, risk prediction for bone fracture, as well as personalized treatment and monitoring of the treatment efficacy.

Hormone therapy is effective for the prevention and treatment of postmenopausal osteoporosis as estrogen reduction is a crucial pathogenic factor. Because of the well-documented side effects, e.g. cardiovascular events and breast cancer risk, estrogen-based therapies are now limited to short-term use ^35, 36^. Another widely used treatment for osteoporosis is bisphosphonates which possess a high affinity for bone minerals with inhibitory effects on osteoclast cells ^36, 37^. However, there is a need for alternative treatments due to the side effects of bisphosphonates, e.g. atypical femur fracture and osteonecrosis of the jaw ^37, 38^. In addition, the treatment effectiveness after 5 years remains uncertain ^39^. A better understanding of the bone remodeling process can help maximize anti-fracture efficacy and minimize adverse skeletal effects.

The clear demonstration of osteoporotic bone loss caused by the disruption of miR-27a suggests its supplementation be explored as a new therapeutic approach. The clinical application requires a system for osteoclast delivery of miR-27a. However, this may be complicated by the negative effects of the miR23a cluster on osteoblast-mediated bone formation ^20^. High levels of miR23a in the mast cells also lead to bone loss through the release of extracellular vesicles by neoplastic mast cells ^40^. Therefore, the targeting miR-27a to the bone resorption surfaces with synthetic compounds such as bisphosphonates or osteoclast-targeting molecules such as acid octapeptides with aspartic acid is critical for future clinical applications ^41^.

## Materials and Methods

### Animals

The CRISPR/Cas9 gene edition strategy was used to generate *ΔmiR-23a∼27a* and *ΔmiR-27a* mouse strains ^42, 43^. Complementary oligonucleotides containing the *miR-23a* or *miR-27a* sgRNA target sequences were designed by CRISPR Design Tool (http://crispr.mit.edu) and inserted into the pX335 plasmid (Addgene, Cambridge, MA), followed by DNA sequencing to verify the correct cloning. A mixture of sgRNA plasmid, Cas9 protein (NEB, Ipswich, MA), and ssODN (Integrated DNA Technologies, Coralville, IA) was injected into the pronuclei and cytoplasm of fertilized eggs ^44, 45^. The survived embryos were transferred into the oviduct of pseudopregnant females for carrying to term. The founder lines were genotyped by PCR analysis using primers 5’- GAC CCA GCC TGG TCA AGA TA-3’ and 5’-GGA CTC CTG TTC CTG CTG AA-3’ to determine the success of the gene edition and germline transmission. Both male and female mice were used in this study. Care and use of experimental animals described in this work comply with guidelines and policies of the University Committee on Animal Resources at the University of Rochester and IACUC at the Forsyth Institute.

### Genes

Total RNAs including miRNAs were isolated using mirVana miRNA isolation kit (Thermo Fisher Scientific, Waltham, MA), followed by polyadenylation using E. coli Poly(A) polymerase (NEB), and reserve transcribed into DNA using Reverse Transcriptase (Thermo Fisher Scientific) and an anchor primer. Based on the dominant mature miRNA sequences, four to six nucleotides were added to the 5’ end to enhance the annealing specificity. To detect the expression of the miRNAs, the reverse transcription products were subject to PCR analysis using forward and reverse primers listed in Supplementary Table 1. The PCR was performed by denaturation at 95^0^C for 2 min and 27 cycles of amplification (95^0^C for 20 s, 60^0^C for 10 s, and 70^0^C for 10 s). The Lentivirus-miR27a (Cat #MmiR3347MR03, GeneCopoeia, Rockville, MD) and Lentivirus-shRNA p62/Sqstm1 (Product ID: MSH093992, Accession: NM_011018.3, GeneCopoeia) were used to express miR-27a and knockdown p62, respectively.

### Cells

Primary cells were harvested from bone marrows of the bilateral femur to obtain bone marrow cells. These cells were incubated with Mouse BD Fc Block™ (Cat #553142, BD Biosciences, San Jose, CA) to reduce non-specific antibody staining caused by receptors for IgG, followed by FACS analysis with anti-CD11b/Mac-1 (M1/70)-APC (17-0112-81, eBioscience/Thermo Fisher Scientific; 1:400), anti-granulocyte mAb: anti-Ly-6G/Gr-1 (RB6-8C5)-PE (12-5931-81, eBioscience; 1:10000), anti-dendritic cell mAb: anti-CD11c (N418)-FITC (11-0114-81, eBioscience; 1:20) using LSR II (BD Biosciences). For OC differentiation, isolated cells were cultured in αMEM containing 10% FBS, 1% L-glutamine, 1% non-essential amino acids, and 5 ng/ml M-CSF for 2 days, followed by the addition of RANKL 10 ng/ml (R&D, Minneapolis, MN). The differentiated cells were then fixed with 10% formalin followed by TRAP staining ^24^.

For bone resorption assay, 5 x 10^4^ primary osteoclast progenitors isolated from 3-month-old control and ΔmiR-27a femur were seeded in 96 well Corning Osteo Assay Surface plate (Cat #3988, Corning, Glendale, AZ), and cultured in 5% CO2 with differentiation media at 37°C. After culturing for 5 days, the resorbed areas on the plates were imaged by a Nikon SMZ1500 microscope, followed by ImageJ analysis. For actin ring formation assay, 5 x 10^4^ primary osteoclast progenitors isolated from 3-month-old control and ΔmiR-27a femur were seeded in 96 well plates. After 5 days of culture, the cells were fixed with 4% PFA in PBS for 15 minutes, permeabilized with 0.1% Triton X-100 in PBS, and incubate with Phalloidin-Alexa 488 (Cat #A12379 Invitrogen) for 1 hour at room temperature. Nuclear counterstaining was performed by DAPI (4’,6-diamidino-2-phenylindole).

### 3’UTR assay

The LUC-3’UTR reporter DNA plasmids contain the 3’UTR from Sqstm1, Tgfbr1, Snx10, Map2k7, and Ctsk fused to the end of a luciferase reporter gene (MmiT079315-MT06, MmiT096143-MT06, MmiT073076-MT06, MmiT099033-MT06, MmiT091679-MT06, GeneCopoeia). C3H10T1/2 mesenchymal cells were transfected by the LUC-3’UTR without or with co-transfection of miR27a (MmiR3347-MR04-50, GeneCopoeia) using Lipofectamine 200 (Cat #11668027, Invitrogen, Waltham, MA).

The transcript stability and its translation efficiency in the presence or absence of miR-27a were determined by the luciferase assay 72 hours after the transfection using the Dual-Luciferase Reporter Assay System (Cat #1910, Promega, Madison, WI). The luminescent intensity was measured by the SpectraMax iD3 Multi-Mode Microplate Reader (Molecular Devices, San Jose, CA). Firefly luciferase activities were normalized by the values of Renilla luciferase.

### Bone analysis and staining

Mouse limbs and spines were collected for ex vivo μCT imaging using a vivaCT40 scanner (Scanco USA, Wayne, PA). The scanned images were segmented for reconstruction to access the relative bone volume (BV/TV, %) by 3-D visualization and analysis using Amira (FEI, Thermo Fisher Scientific). To perform quantitative measurements, 170 and 200 slices were used for the femur trabecular bone and L5 spine, respectively. Cortical bone thickness (Ct. Th, mm) was analyzed at a standardized location of 30 slices near the midshaft. Skeletal preparation, fixation, and embedding for paraffin sections were performed as described ^46–55^. Samples were subject to hematoxylin/eosin staining for histology, TRAP staining, van Kossa staining, or immunological staining with avidin: biotinylated enzyme complex ^46, 47, 49, 51, 52, 54–59^. The immunological staining was visualized by enzymatic color reaction or fluorescence according to the manufacturer’s specification (Vector Laboratories, Burlingame, CA). After TRAP staining, osteoclast number per bone area (N.Oc/T.Ar), osteoclast number per bone surface (N.Oc/BS), and osteoclast surface per bone surface (Oc.S/BS, %) was determined for statistical significance. Rabbit antibodies Collagen I (LSL-LB-1190, Cosmo Bio Co., LTD., Japan; 1:2000), Cathepsin K (ab19027, Abcam, Cambridge, MA; 1:50); mouse antibodies Osteopontin (MPIIIB10, Hybridoma Bank, Iowa; 1:500) were used in these analyses. Images were taken using Leica DM2500 and DFC310FX imaging system (Leica, Bannockburn, IL) and Zeiss Axio Observer microscope (Carl Zeiss, Thornwood, NY). Bone formation rate (BFR) was examined by double labeling of Alizarin Red S and Calcein Green, injected intraperitoneally with a 7-day interval. The labeled samples were embedded without decalcification for frozen sections (SECTION-LAB Co. Ltd, Japan) ^49, 54, 55^, followed by analyzed under a fluorescent microscope using an OsteoMeasure morphometry system (OsteoMetrics, Atlanta, GA) to determine Mineral Apposition Rate (MAR, BFR/BS).

### Statistics and Reproducibility

Microsoft Excel 2010 was used for statistical analysis. The significance was determined by two-sided student t-tests. A *p*-value less than 0.05 was considered statistically significant. Before performing the t-tests, the normality of the data distribution was first validated by the Shapiro-Wilk normality test. Analysis of samples by µCT was performed by a technician who is blinded to the condition. No randomization, statistical method to predetermine the sample size, and inclusion/exclusion criteria defining criteria for samples were used. At least 3 independent experiments were performed for statistical analyses of the animal tissues described in figure legends. Statistical data were presented as mean ± SD. The miPath v.3 KEGG Reverse Search (https://dianalab.e-ce.uth.gr/html/mirpathv3/index.php?r=mirpath/reverse) with Search Pathway: mmu04380 and Method: TarBase v7.0 was used to identify the candidates that are both miR-27a targets (TarBase v7.0) and genes associated with osteoclast differentiation (KEGG: mmu04380).

### Materials & Correspondence

Correspondence and material requests should be addressed to W.H. and T.M.,

## ACKNOWLEDGEMENTS

We thank Chyuan-Sheng Lin, Keiko Kaneko, Ya-Hui Chiu, Michael Thullen, and Hsiao-Man Ivy Yu for assistance in transgenic mouse strains, plasmid DNA construction and purification, FACS, µCT scanning, and imaging analysis, respectively. Research reported in this publication is supported by the National Institute of Dental and Craniofacial Research of the National Institutes of Health under award numbers R01DE015654 and R01DE026936 to W.H. and R21DE028696 to T.M.

## Supplementary Information for

**Fig. S1.**
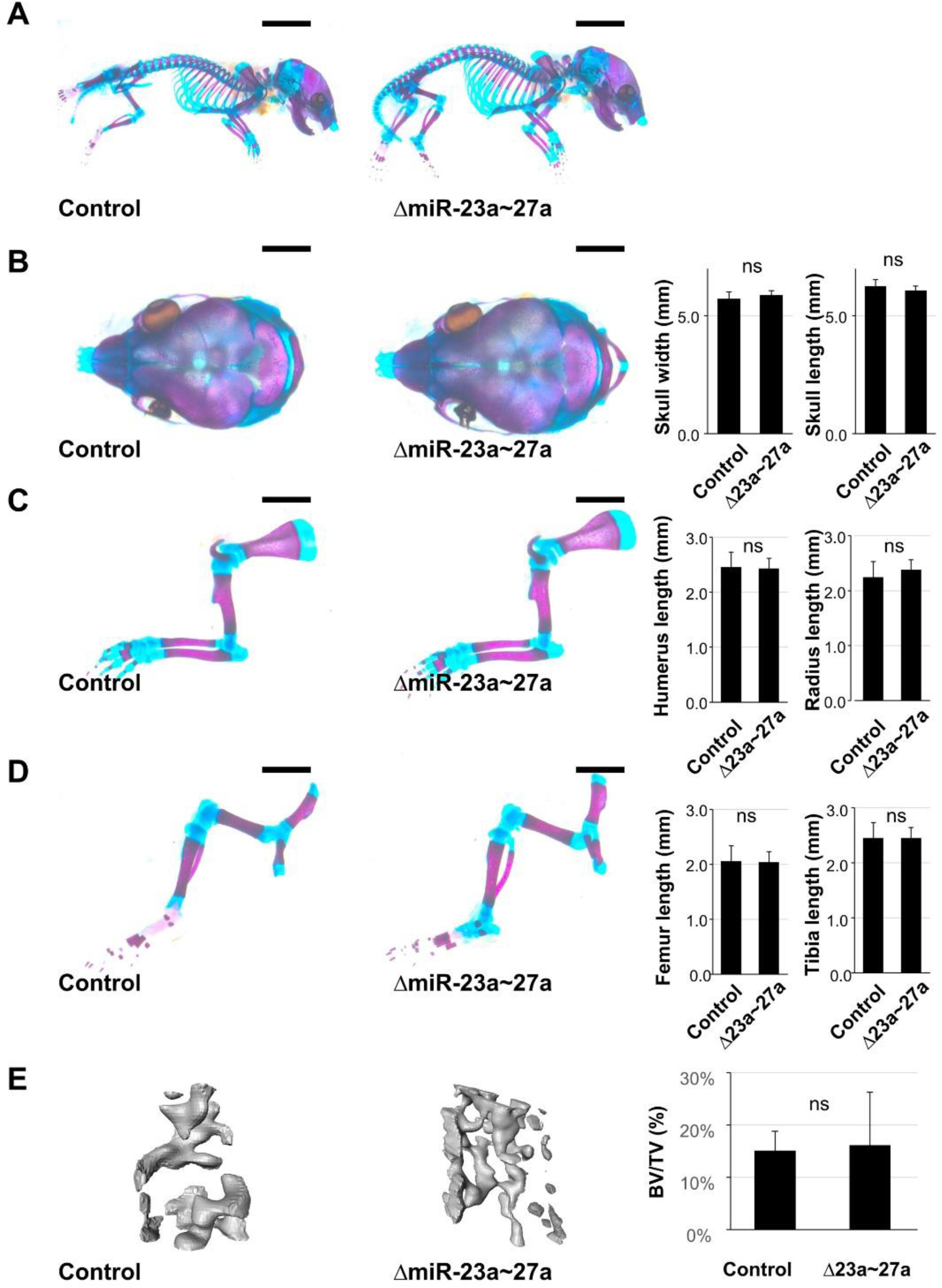
MiR-23a and miR-27a are dispensable during skeletal development. (A-D) Skeletal staining of control (n=9) and ΔmiR-23a∼27a (n=8) mice reveals no skeletal deformities in the skull (A and B), forelimb (A and C), hindlimb (A and D) at postnatal day 0 (P0). The statistical evaluation shows no significant difference between the control and mutant in the skull width (B; n=8, mean ± SD mm, p-value > 0.24, two-sided student t-test) and length (B; n=8, mean ± SD mm, *P* > 0.16), in the humerus (C; n=8, mean ± SD mm, p-value > 0.77) and radius (C; n=8, mean ± SD mm, p-value > 0.28), in the femur (D; n=8, mean ± SD mm, p-value > 0.76) and tibia (D; n=8, mean ± SD mm, p-value > 0.99). (E) Micro-computed tomography (µCT) analysis of femur trabecular bone volume shows no significant changes in ΔmiR-23a∼27a mice compared to the control (n=8, mean ± SD %, p-value > 0.77, two-sided student t-test). Scale bars, 5 mm (A); 2 mm (B-D).

**Fig. S2.**
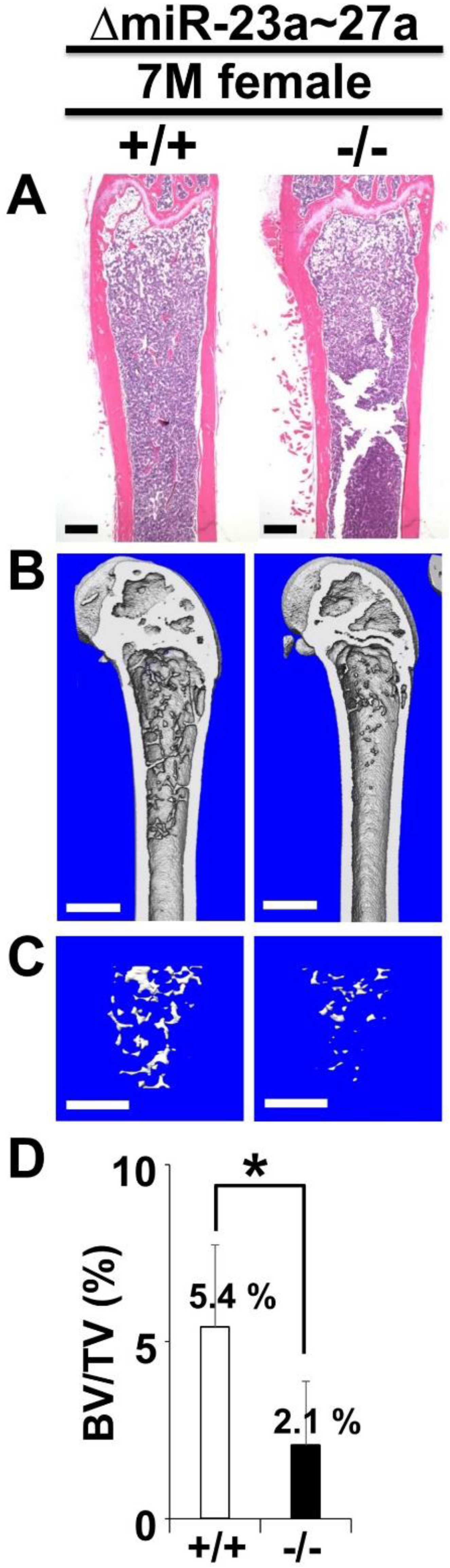
The loss of miR-23a∼27a causes an osteopenic phenotype in mice. Femurs of the 7-month-old (7M) wild-type (+/+) and mutant (-/-) females, were analyzed by μCT scanning, followed by H&E staining (A). The 3D rendering μCT images of the distal femur (B) and femoral metaphysis (C) were subject to quantitative analysis (D) for BV/TV (trabecular bone volume per tissue volume, n=3, *, p-value < 0.05, mean ± SD; student t-test). Scale bars, 500 µm (A-C).

**Fig. S3.**
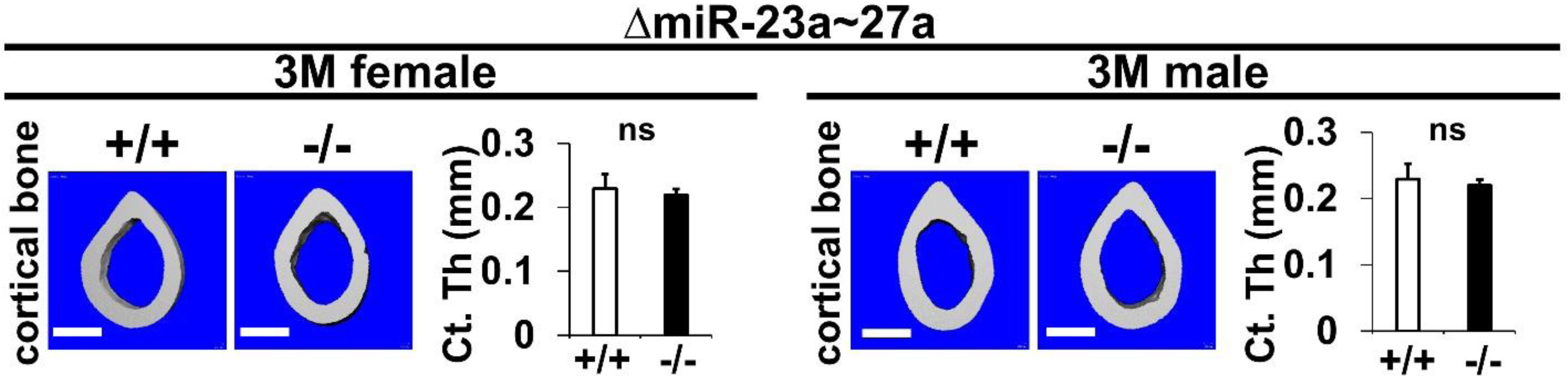
The loss of miR-23a∼27a does not affect cortical bone thickness. Femurs of the 3-month-old (3M) ΔmiR-23a∼27a males and females were analyzed by μCT scanning. Images show the reconstructed cortical bone of wild-type (+/+) and mutant (-/-). Quantitative analyses of cortical (Ct.) thickness (Th) are shown in graphs (n=5, mean ± SD; student t-test, ns, non-significant). Images are representatives of five independent experiments. Scale bars, 500 µm.

**Fig. S4.**
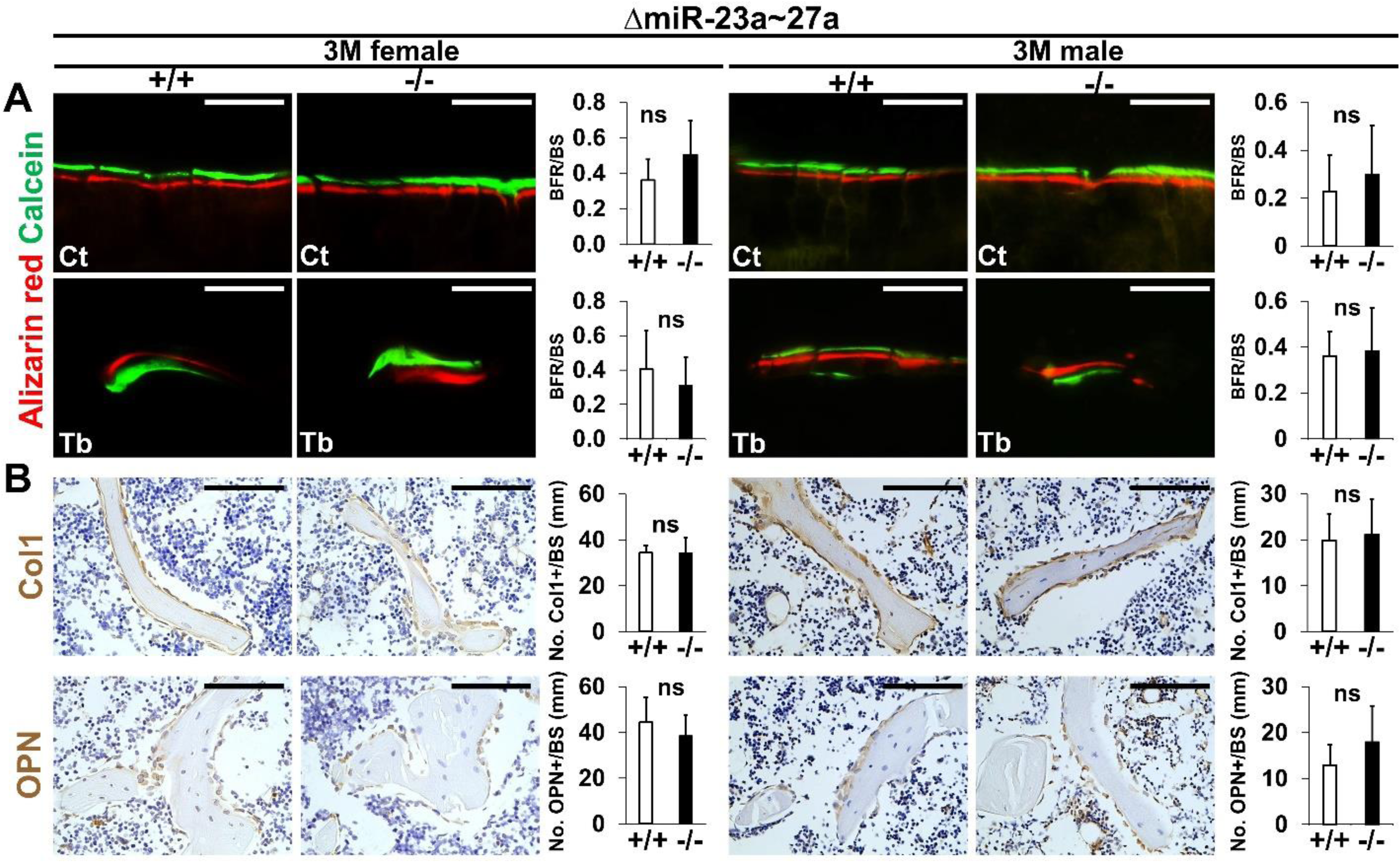
Bone formation and osteoblastogenesis are not affected by the ΔmiR-23a∼27a deletion. Sections of the 3-month-old (3M) ΔmiR-23a∼27a males and females were analyzed by (A) double labeling of alizarin red and calcein, and (B) immunostaining of type 1 collagen (Col1) and Osteopontin (OPN). Quantitative analyses of (A) bone formation rate per bone surface (BFR/BS, n=3, mean ± SD; student t-test) and (B) number of positively stained cells over the bone surface (No. +/BS, n=5, mean ± SD; student t-test, ns, non-significant) in the wild-type (+/+) and mutant (-/-) distal femurs are shown in graphs. Eb, endosteal; Tb, trabecular. Images are representatives of three independent experiments. Scale bar, 100 µm.

**Fig. S5.**
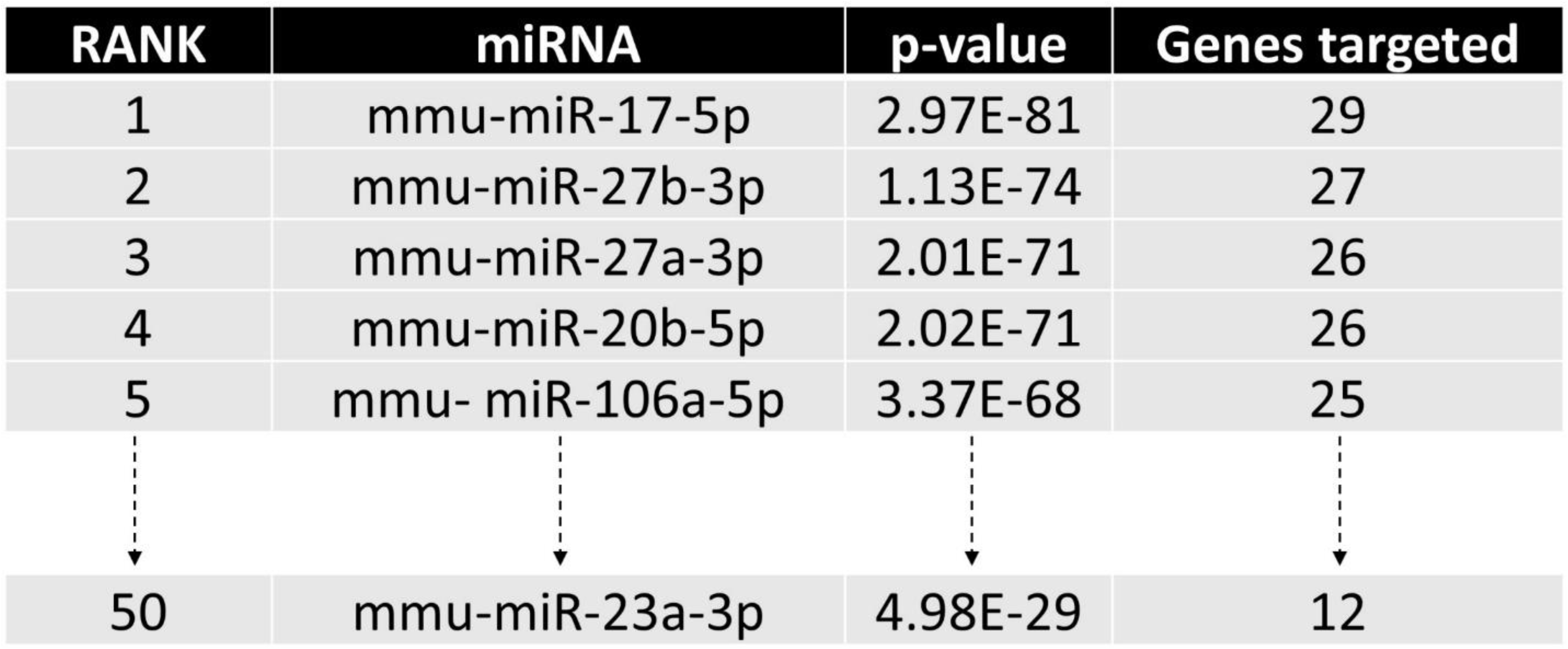
Identification of miRNA candidates in osteoclast differentiation pathway. The diagram illustrates the top candidates identified by the miRPath Reverse-Search module based on the target accumulation in osteoclast differentiation-related genes in the Kyoto Encyclopedia of Genes ad Genomics (KEGG: mmu04380).

**Fig. S6.**
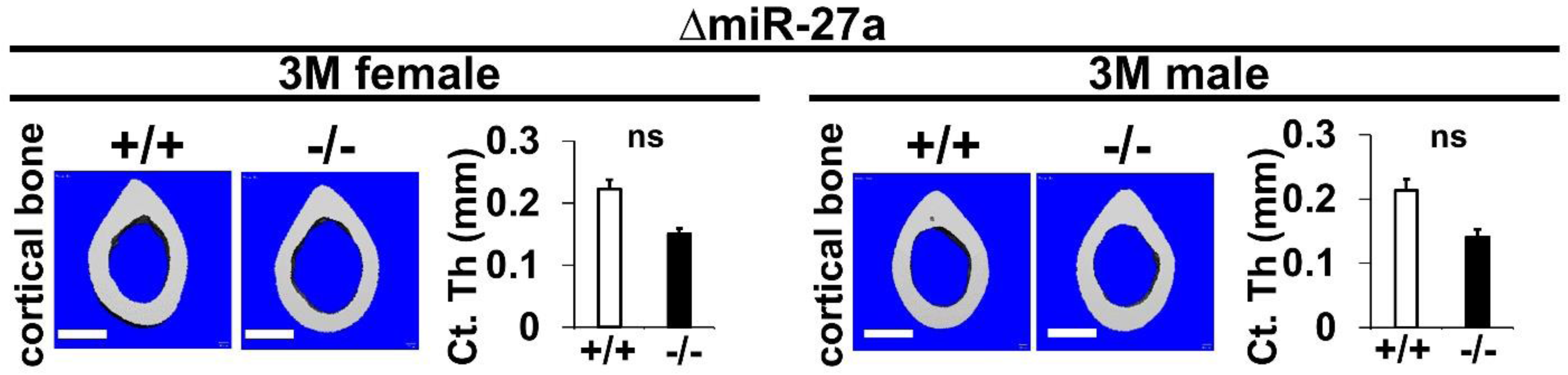
The loss of miR-27a does not affect cortical bone thickness. Femurs of the 3-month-old (3M) and ΔmiR-27a males and females were analyzed by μCT scanning. Images show the reconstructed cortical bone of wild-type (+/+) and mutant (-/-). Quantitative analyses of cortical (Ct.) thickness (Th) are shown in graphs (n=5, mean ± SD; student t-test, ns, non-significant). Images are representatives of five independent experiments. Scale bars, 500 µm.

**Fig. S7.**
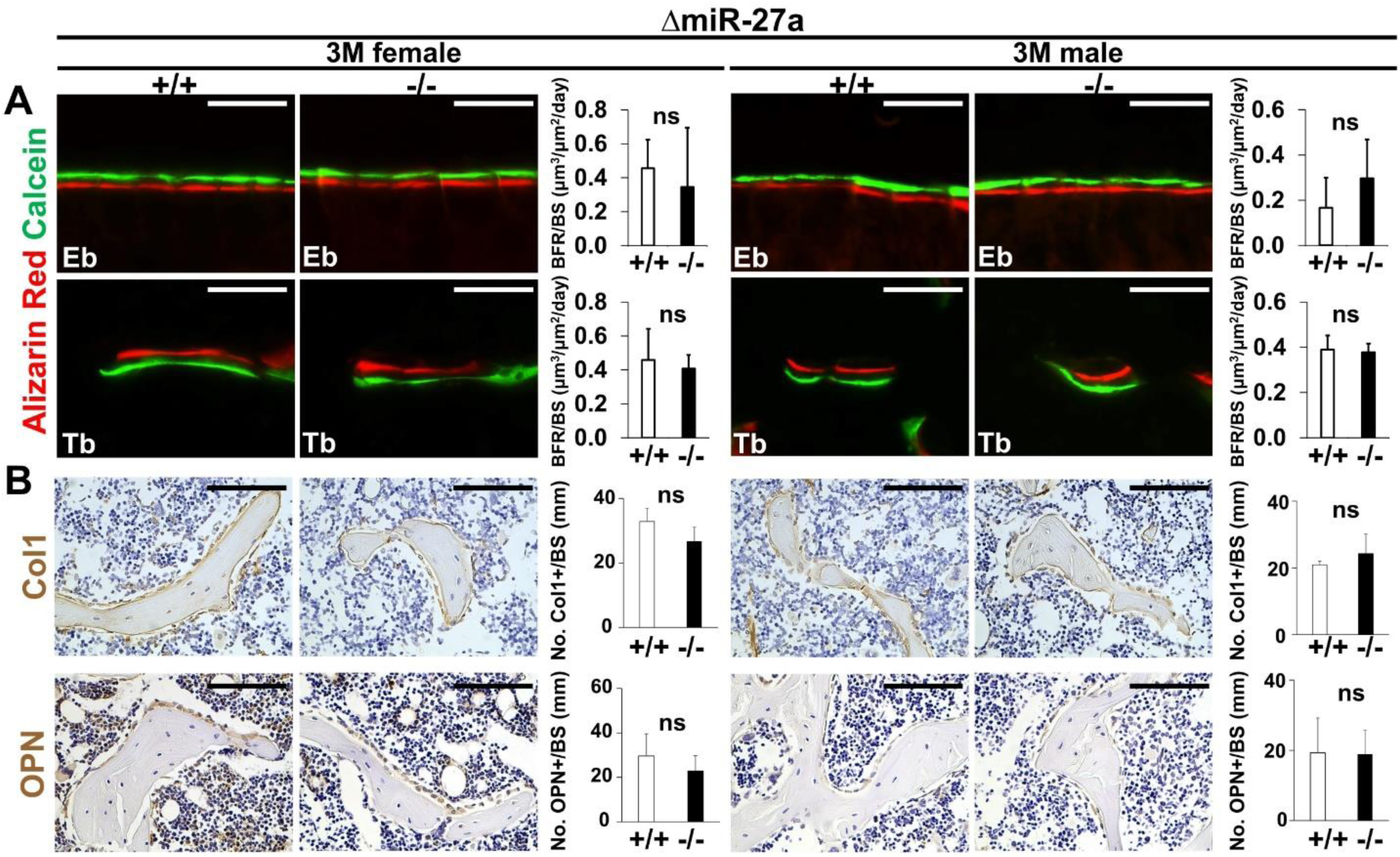
Bone formation and osteoblastogenesis are not affected by the ΔmiR-27a deletion. Sections of the 3-month-old (3M) ΔmiR-27a males and females were analyzed by (A) double labeling of alizarin red and calcein, and (B) immunostaining of type 1 collagen (Col1) and Osteopontin (OPN). Quantitative analyses of (A) bone formation rate per bone surface (BFR/BS, n=3, mean ± SD; student t-test) and (B) number of positively stained cells over the bone surface (No. +/BS, n=5, mean ± SD; student t-test, ns, non-significant) in the wild-type (+/+) and mutant (-/-) distal femurs are shown in graphs. Eb, endosteal; Tb, trabecular. Images are representatives of three independent experiments. Scale bar, 100 µm.

**Fig. S8.**
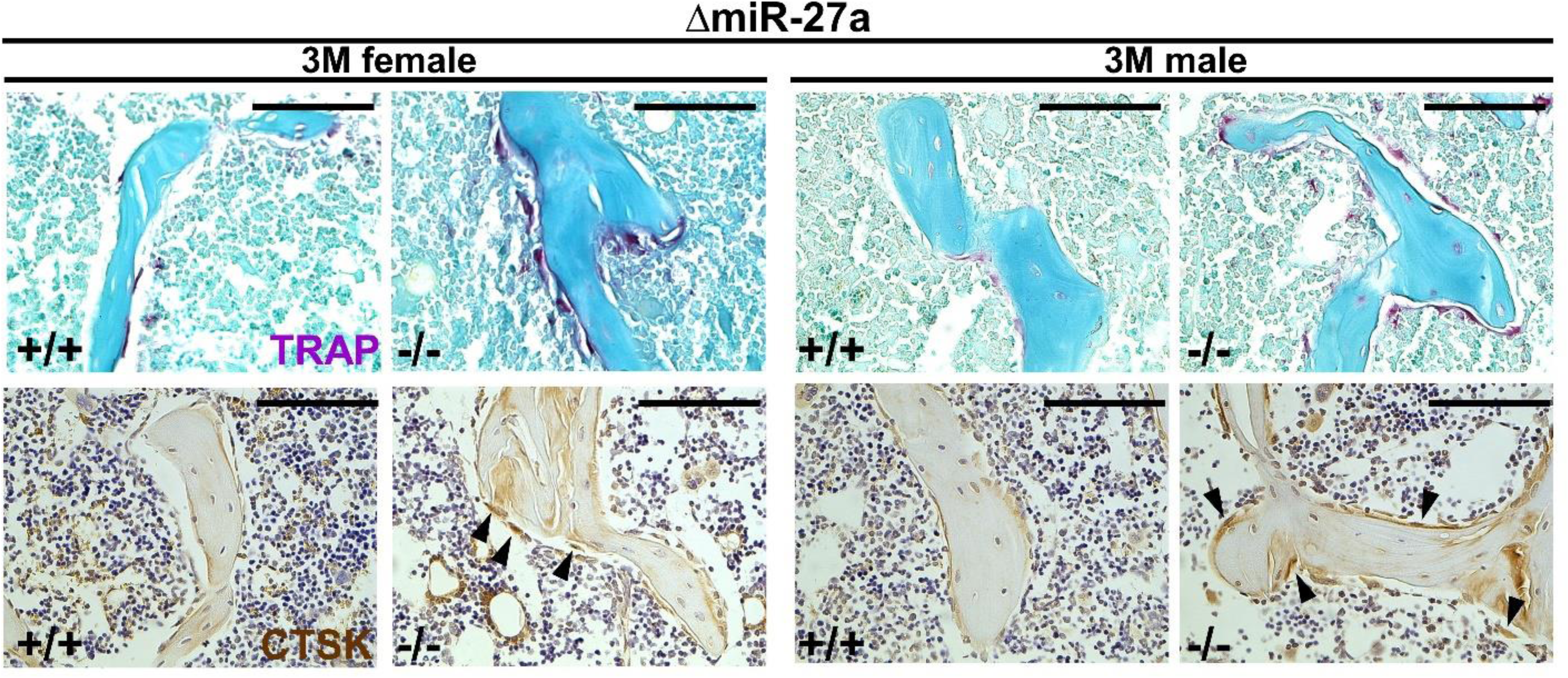
Enhanced osteoclastogenesis in the ΔmiR-27a mice. Sections of the 3-month-old (3M) ΔmiR-27a males and females were analyzed by tartrate-resistant acid phosphatase (TRAP) staining and immunostaining of Cathepsin K (CTSK). Based on the positively stained cells in the wild-type (+/+) and mutant (-/-) distal femurs (No. +/BS, n=5, mean ± SD; student t-test), quantitative analyses were performed to obtain histomorphometric parameters of bone resorption, the number of osteoclast/bone area (N.Oc/T.Ar), osteoclast surface/bone surface (Oc.S/BS), Osteoclast number/bone surface (N.Oc/BS) shown in Fig. 5. Arrowheads indicate differentiated osteoclast cells positive for TRAP and CTSK. Images are representatives of five independent experiments. Scale bars, 100 µm.

**Fig. S9.**
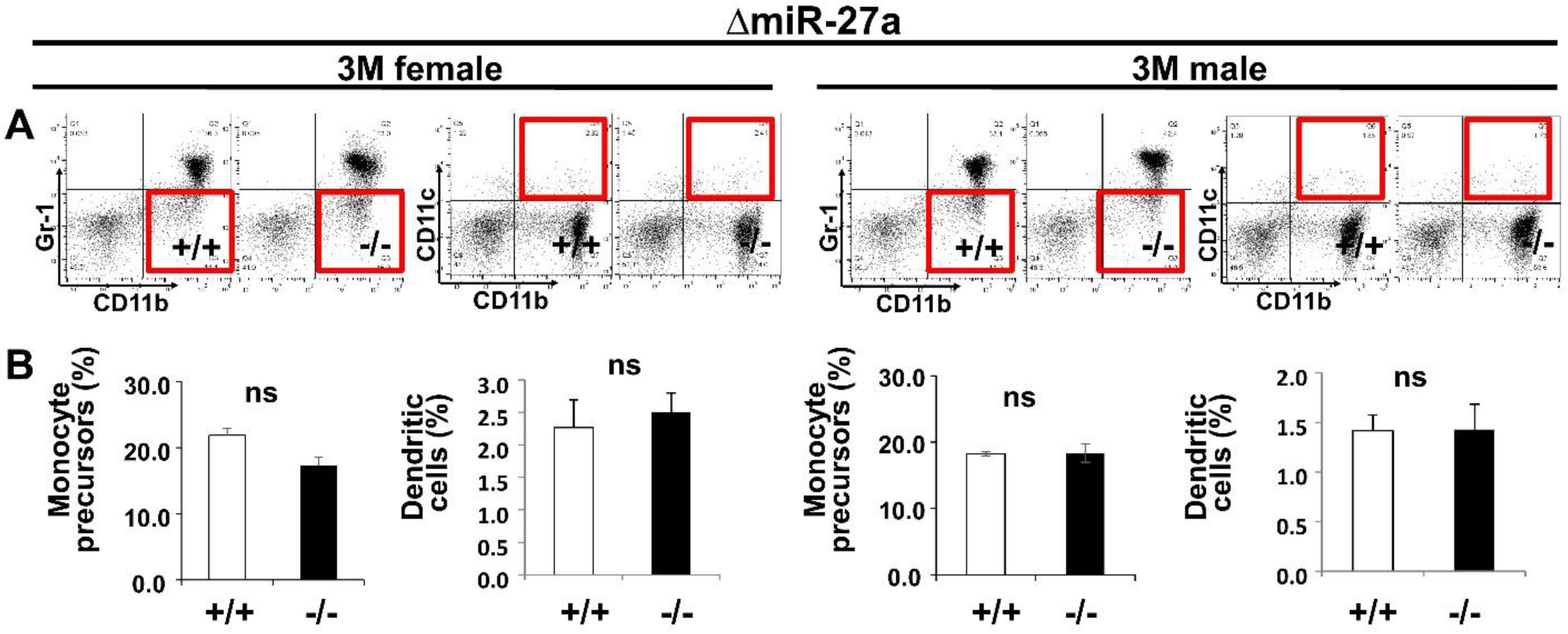
Osteoclast precursor populations are not affected by the loss of miR-27a. (A) FACS analysis examines the CD11b+/Gr-1– and CD11b+/CD11c+ populations for monocyte precursors and dendritic cells, respectively. Images are representatives of three independent experiments. (B) No significant difference in the bone marrow of 3-month-old (3M) wild-type (+/+) and ΔmiR-23a∼27a (-/-) mice (n = 3, mean ± SD; student t-test).

**Fig. S10.**
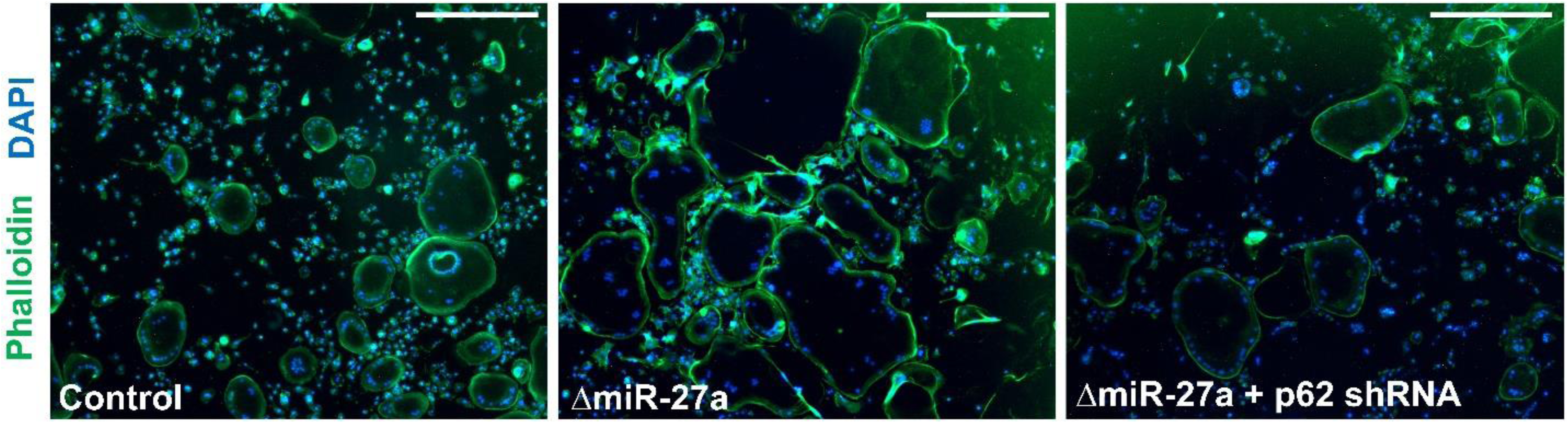
No effect of actin ring formation by the loss of miR-27a in osteoclast cells. The actin ring formation in the OC cells of control, ΔmiR-27a, and ΔmiR-27a plus shRNA-mediated p62 knockdown is examined by phalloidin staining after 5 days of differentiation with RANKL and MCSF. Scale bars, 500 µm.

**Table S1.**
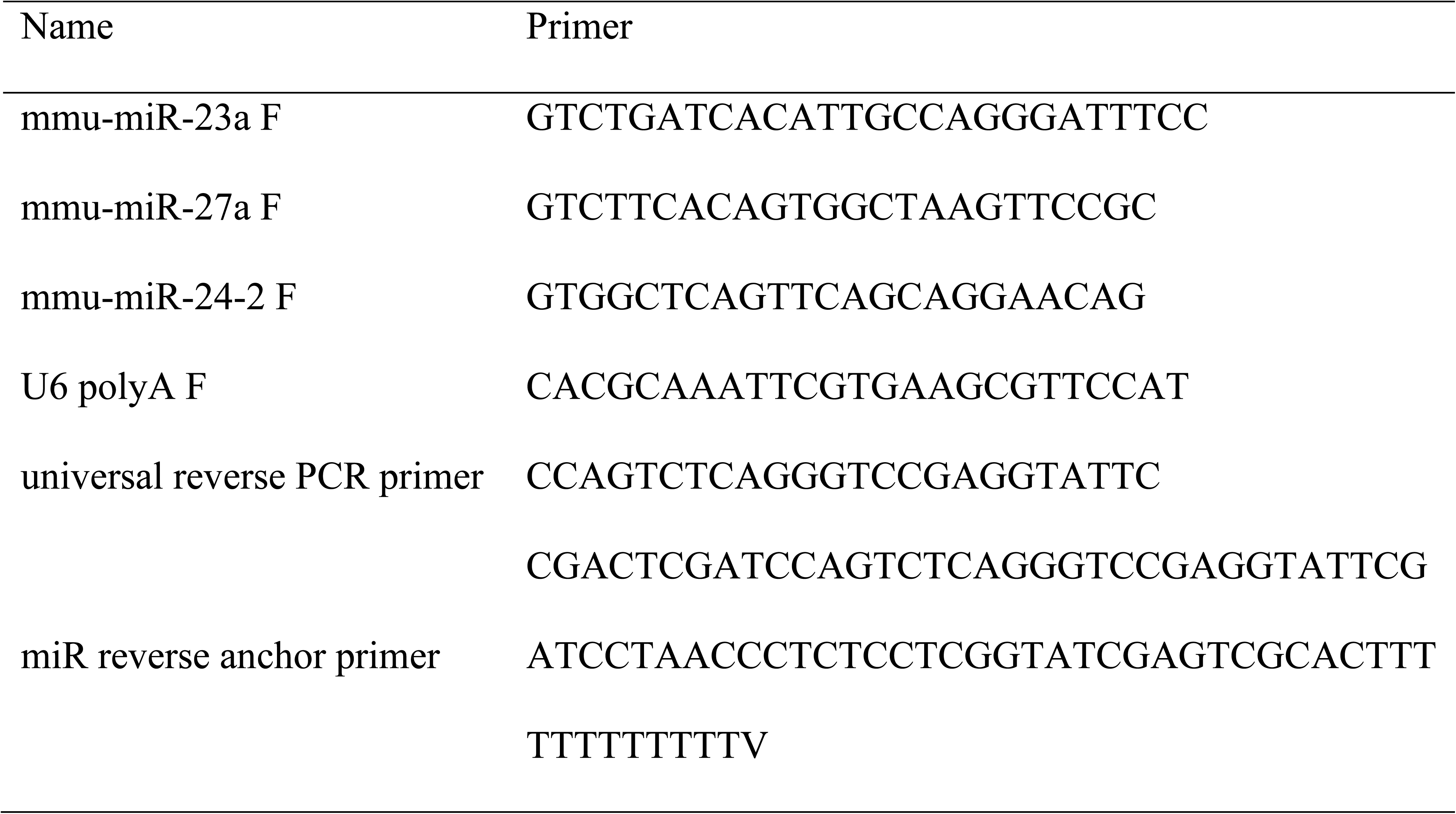
Primers for RT-PCR analysis of miR23a∼27a∼24-2.

## References

1. Bartel DP. MicroRNAs: genomics, biogenesis, mechanism, and function. Cell 116, 281–297 (2004).

2. Baek D, Villen J, Shin C, Camargo FD, Gygi SP, Bartel DP. The impact of microRNAs on protein output. Nature 455, 64–71 (2008).

3. Hausser J, Zavolan M. Identification and consequences of miRNA-target interactions--beyond repression of gene expression. Nat Rev Genet 15, 599–612 (2014).

4. Stark A, Brennecke J, Bushati N, Russell RB, Cohen SM. Animal MicroRNAs confer robustness to gene expression and have a significant impact on 3’UTR evolution. Cell 123, 1133–1146 (2005).

5. Ebert MS, Sharp PA. Roles for microRNAs in conferring robustness to biological processes. Cell 149, 515–524 (2012).

6. Li Z, Rana TM. Therapeutic targeting of microRNAs: current status and future challenges. Nat Rev Drug Discov 13, 622–638 (2014).

7. Miska EA, et al. Most Caenorhabditis elegans microRNAs are individually not essential for development or viability. PLoS genetics 3, e215 (2007).

8. Park CY, et al. A resource for the conditional ablation of microRNAs in the mouse. Cell reports 1, 385–391 (2012).

9. Croce CM. Causes and consequences of microRNA dysregulation in cancer. Nat Rev Genet 10, 704–714 (2009).

10. Shenoy A, Blelloch RH. Regulation of microRNA function in somatic stem cell proliferation and differentiation. Nature reviews 15, 565–576 (2014).

11. Manolagas SC, Jilka RL. Bone marrow, cytokines, and bone remodeling. Emerging insights into the pathophysiology of osteoporosis. N Engl J Med 332, 305–311 (1995).

12. Feng X, McDonald JM. Disorders of bone remodeling. Annu Rev Pathol 6, 121–145 (2011).

13. Gaur T, et al. Dicer inactivation in osteoprogenitor cells compromises fetal survival and bone formation, while excision in differentiated osteoblasts increases bone mass in the adult mouse. Developmental biology 340, 10–21 (2010).

14. Mizoguchi F, et al. Osteoclast-specific Dicer gene deficiency suppresses osteoclastic bone resorption. Journal of cellular biochemistry 109, 866–875 (2010).

15. Seeliger C, et al. Five freely circulating miRNAs and bone tissue miRNAs are associated with osteoporotic fractures. J Bone Miner Res 29, 1718–1728 (2014).

16. Guo Q, Chen Y, Guo L, Jiang T, Lin Z. miR-23a/b regulates the balance between osteoblast and adipocyte differentiation in bone marrow mesenchymal stem cells. Bone Res 4, 16022 (2016).

17. You L, Pan L, Chen L, Gu W, Chen J. MiR-27a is Essential for the Shift from Osteogenic Differentiation to Adipogenic Differentiation of Mesenchymal Stem Cells in Postmenopausal Osteoporosis. Cell Physiol Biochem 39, 253–265 (2016).

18. Hassan MQ, et al. A network connecting Runx2, SATB2, and the miR-23a∼27a∼24-2 cluster regulates the osteoblast differentiation program. Proceedings of the National Academy of Sciences of the United States of America 107, 19879–19884 (2010).

19. Guo L, et al. Estrogen inhibits osteoclasts formation and bone resorption via microRNA-27a targeting PPARgamma and APC. J Cell Physiol 234, 581–594 (2018).

20. Zeng HC, et al. MicroRNA miR-23a cluster promotes osteocyte differentiation by regulating TGF-beta signalling in osteoblasts. Nature communications 8, 15000 (2017).

21. Androsavich JR, et al. Polysome shift assay for direct measurement of miRNA inhibition by anti-miRNA drugs. Nucleic acids research 44, e13 (2016).

22. Ebert MS, Neilson JR, Sharp PA. MicroRNA sponges: competitive inhibitors of small RNAs in mammalian cells. Nature methods 4, 721–726 (2007).

23. Charles JF, Nakamura MC. Bone and the innate immune system. Curr Osteoporos Rep 12, 1–8 (2014).

24. Yao Z, et al. Tumor necrosis factor-alpha increases circulating osteoclast precursor numbers by promoting their proliferation and differentiation in the bone marrow through up-regulation of c-Fms expression. J Biol Chem 281, 11846–11855 (2006).

25. Kanehisa M, Goto S. KEGG: kyoto encyclopedia of genes and genomes. Nucleic acids research 28, 27–30 (2000).

26. Kanehisa M, et al. KEGG for linking genomes to life and the environment. Nucleic acids research 36, D480–484 (2008).

27. Vlachos IS, et al. DIANA-TarBase v7.0: indexing more than half a million experimentally supported miRNA:mRNA interactions. Nucleic acids research 43, D153–159 (2015).

28. Vlachos IS, et al. DIANA-miRPath v3.0: deciphering microRNA function with experimental support. Nucleic acids research 43, W460–466 (2015).

29. Rea SL, et al. A novel mutation (K378X) in the sequestosome 1 gene associated with increased NF-kappaB signaling and Paget’s disease of bone with a severe phenotype. J Bone Miner Res 21, 1136–1145 (2006).

30. Duran A, et al. The atypical PKC-interacting protein p62 is an important mediator of RANK-activated osteoclastogenesis. Developmental cell 6, 303–309 (2004).

31. Ma Y, et al. Validation of downregulated microRNAs during osteoclast formation and osteoporosis progression. Mol Med Rep 13, 2273–2280 (2016).

32. Ou M, et al. Identification of potential microRNA-target pairs associated with osteopetrosis by deep sequencing, iTRAQ proteomics and bioinformatics. Eur J Hum Genet 22, 625–632 (2014).

33. Laurin N, Brown JP, Morissette J, Raymond V. Recurrent mutation of the gene encoding sequestosome 1 (SQSTM1/p62) in Paget disease of bone. Am J Hum Genet 70, 1582–1588 (2002).

34. Garnero P. Biomarkers for osteoporosis management: utility in diagnosis, fracture risk prediction and therapy monitoring. Mol Diagn Ther 12, 157–170 (2008).

35. Rossouw JE, et al. Risks and benefits of estrogen plus progestin in healthy postmenopausal women: principal results From the Women’s Health Initiative randomized controlled trial. Jama 288, 321–333 (2002).

36. Khosla S, Hofbauer LC. Osteoporosis treatment: recent developments and ongoing challenges. Lancet Diabetes Endocrinol 5, 898–907 (2017).

37. Russell RG, Watts NB, Ebetino FH, Rogers MJ. Mechanisms of action of bisphosphonates: similarities and differences and their potential influence on clinical efficacy. Osteoporos Int 19, 733–759 (2008).

38. Shane E, et al. Atypical subtrochanteric and diaphyseal femoral fractures: second report of a task force of the American Society for Bone and Mineral Research. J Bone Miner Res 29, 1–23 (2014).

39. Izano MA, et al. Bisphosphonate Treatment Beyond 5 Years and Hip Fracture Risk in Older Women. JAMA Netw Open 3, e2025190 (2020).

40. Kim DK, et al. Mastocytosis-derived extracellular vesicles deliver miR-23a and miR-30a into pre-osteoblasts and prevent osteoblastogenesis and bone formation. Nature communications 12, 2527 (2021).

41. Dang L, et al. Targeted Delivery Systems for Molecular Therapy in Skeletal Disorders. International journal of molecular sciences 17, 428 (2016).

42. Cong L, et al. Multiplex genome engineering using CRISPR/Cas systems. Science (New York, NY 339, 819–823 (2013).

43. Mali P, et al. RNA-guided human genome engineering via Cas9. Science (New York, NY 339, 823–826 (2013).

44. Mashiko D, Fujihara Y, Satouh Y, Miyata H, Isotani A, Ikawa M. Generation of mutant mice by pronuclear injection of circular plasmid expressing Cas9 and single guided RNA. Scientific reports 3, 3355 (2013).

45. Wu WH, et al. CRISPR Repair Reveals Causative Mutation in a Preclinical Model of Retinitis Pigmentosa. Molecular therapy : the journal of the American Society of Gene Therapy 24, 1388–1394 (2016).

46. Maruyama T, Mirando AJ, Deng CX, Hsu W. The balance of WNT and FGF signaling influences mesenchymal stem cell fate during skeletal development. Sci Signal 3, ra40 (2010).

47. Yu HM, et al. The role of Axin2 in calvarial morphogenesis and craniosynostosis. Development (Cambridge, England) 132, 1995–2005 (2005).

48. Mirando AJ, Maruyama T, Fu J, Yu HM, Hsu W. Beta-catenin/cyclin D1 mediated development of suture mesenchyme in calvarial morphogenesis. BMC Dev Biol 10, 116 (2010).

49. Maruyama T, Jiang M, Hsu W. Gpr177, a novel locus for bone mineral density and osteoporosis, regulates osteogenesis and chondrogenesis in skeletal development. J Bone Miner Res 28, 1150–1159 (2013).

50. Yu HM, Jin Y, Fu J, Hsu W. Expression of Gpr177, a Wnt trafficking regulator, in mouse embryogenesis. Dev Dyn 239, 2102–2109 (2010).

51. Yu HM, Liu B, Chiu SY, Costantini F, Hsu W. Development of a unique system for spatiotemporal and lineage-specific gene expression in mice. Proceedings of the National Academy of Sciences of the United States of America 102, 8615–8620 (2005).

52. Yu HM, Liu B, Costantini F, Hsu W. Impaired neural development caused by inducible expression of Axin in transgenic mice. Mechanisms of development 124, 146–156 (2007).

53. Maruyama EO, Yu HM, Jiang M, Fu J, Hsu W. Gpr177 deficiency impairs mammary development and prohibits Wnt-induced tumorigenesis. PLoS ONE 8, e56644 (2013).

54. Maruyama T, Jeong J, Sheu TJ, Hsu W. Stem cells of the suture mesenchyme in craniofacial bone development, repair and regeneration. Nature communications 7, 10526 (2016).

55. Maruyama T, et al. Rap1b is an effector of Axin2 regulating crosstalk of signaling pathways during skeletal development. J Bone Miner Res, (2017).

56. Chiu SY, Asai N, Costantini F, Hsu W. SUMO-Specific Protease 2 Is Essential for Modulating p53-Mdm2 in Development of Trophoblast Stem Cell Niches and Lineages. PLoS biology 6, e310 (2008).

57. Fu J, Hsu W. Epidermal Wnt controls hair follicle induction by orchestrating dynamic signaling crosstalk between the epidermis and dermis. J Invest Dermatol 133, 890–898 (2013).

58. Russell HK, Jr. A modification of Movat’s pentachrome stain. Archives of pathology 94, 187–191 (1972).

59. Fu J, et al. Disruption of SUMO-Specific Protease 2 Induces Mitochondria Mediated Neurodegeneration. PLoS genetics 10, e1004579 (2014).

